# A highly contiguous genome assembly of red perilla (*Perilla frutescens*) domesticated in Japan

**DOI:** 10.1101/2022.09.16.508052

**Authors:** Keita Tamura, Mika Sakamoto, Yasuhiro Tanizawa, Takako Mochizuki, Shuji Matsushita, Yoshihiro Kato, Takeshi Ishikawa, Keisuke Okuhara, Yasukazu Nakamura, Hidemasa Bono

**Affiliations:** Laboratory of Genome Informatics, Graduate School of Integrated Sciences for Life, Hiroshima University, 3-10-23 Kagamiyama, Higashi-Hiroshima, Hiroshima 739-0046, Japan; Laboratory of BioDX, Genome Editing Innovation Center, Hiroshima University, 3-10-23 Kagamiyama, Higashi-Hiroshima, Hiroshima 739-0046, Japan; Genome Informatics Laboratory, Department of Informatics, National Institute of Genetics, 1111 Yata, Mishima, Shizuoka 411-8540, Japan; Agricultural Technology Research Center, Hiroshima Prefectural Technology Research Institute, 6869 Hara, Hachihonmatsucho, Higashi-Hiroshima, Hiroshima 739-0151, Japan; Mishima Foods Co., Ltd., 3-10-7 Kanonshinmachi, Nishi-ku, Hiroshima City, Hiroshima 733-0036, Japan; PtBio Inc., 3-10-23 Kagamiyama, Higashi-Hiroshima, Hiroshima 739-0046, Japan

**Keywords:** *Perilla frutescens*, genome assembly, PacBio HiFi reads

## Abstract

*Perilla frutescens* (Lamiaceae) is an important herbal plant with hundreds of bioactive chemicals, among which perillaldehyde and rosmarinic acid are the two major bioactive compounds in the plant. The leaves of red perilla are used as traditional Kampo medicine or food ingredients. However, the medicinal and nutritional uses of this plant could be improved by enhancing the production of valuable metabolites through the manipulation of key enzymes or regulatory genes using genome editing technology. Here, we generated a high-quality genome assembly of red perilla domesticated in Japan. A near-complete chromosome level assembly of *P. frutescens* was generated contigs with N50 of 41.5 Mb from PacBio HiFi reads. 99.2% of the assembly was anchored into 20 pseudochromosomes, among which seven pseudochromosomes consisted of one contig, while the rest consisted of less than six contigs. Gene annotation and prediction of the sequences successfully predicted 86,258 gene models, including 76,825 protein-coding genes. Further analysis showed that potential targets of genome editing for the engineering of anthocyanin pathways in *P. frutescens* are located on the late-stage pathways. Overall, our genome assembly could serve as a valuable reference for selecting target genes for genome editing of *P. frutescens*.

## 1. Introduction

*Perilla frutescens* is an annual herbal plant belonging to the Lamiaceae family, and it is widely cultivated in Asian countries.^1^ *P. frutescens* is an allotetraploid (2*n* = 4*x* = 40) species, and *P. citriodora* (2*n* = 20) is believed to be one of the diploid genome donors.^2^ There are two chemotypes of perilla plants based on the content of anthocyanins: red and green perilla. Red perilla (“aka-shiso” in Japanese) is an anthocyanin-rich variety with dark red or purple leaves and stems, while green perilla (“ao-jiso” in Japanese) is an anthocyanin deficient variety with green leaves and stems.^3^ Both green and red perilla leaves are often used as a material for cooking. Particularly, the leaves of red perilla are used as traditional Kampo medicine “soyou” to treat stomach problems,^1,2,4,5^ and the seeds are also used to produce oil. Perilla seed oil is a rich source of α-linolenic acid, and its potential health benefits have been reported.^6,7^

Thus far, hundreds of bioactive compounds have been identified in *P. frutescens*,^1,8^ among which perillaldehyde (monoterpenenoid) and rosmarinic acid (phenylpropanoid) are major phytochemicals.^9^ Perillaldehyde has been shown to possess anti-inflammatory,^10^ antidepressant,^11^ antifungal, and antibacterial activities;^12,13^ additionally, rosmarinic acid possesses antiviral, antibacterial, and anti-inflammatory activities.^14^ Several enzymes for the biosynthesis of these compounds have been identified in *P. frutescens*. Perillaldehyde appears to be biosynthesized by the hydroxylation and subsequent oxidation of the C-7 position of limonene. Limonene synthase and a cytochrome P450 monooxygenase (P450) catalyzing the two-step oxidation of limonene at C-7 position have been cloned and characterized in *P. frutescens*.^15,16^ Rosmarinic acid is proposed to be biosynthesized from 4-coumaroyl-CoA and 4-hydroxyphenyllactic acid.^17^ Rosmarinic acid synthase, which is the first specific enzyme for rosmarinic acid biosynthesis, catalyzes the ester formation step of these two compounds. After 4-coumaroyl-4’-hydroxyphenyllactic acid formation, P450 enzymes belonging to the CYP98A family member are known to catalyze the final hydroxylation steps leading to rosmarinic acid production.^14,17^ These enzymes have been cloned and characterized from several plant species, including *Coleus scutellarioides* (Lamiaceae); however, none has been identified in perilla plants.

Recently, genome editing tools, such as CRISPR-Cas9, have been used for the engineering of plant biosynthetic pathways.^18^ For instance, deletion of the autoinhibitory domain of glutamate decarboxylase 3 (*GAD3*) using CRISPR-Cas9 technology promoted the accumulation of GABA (γ-aminobutyric acid) in tomato fruit, and this product is already commercially available.^19,20^ Additionally, silencing of the potato sterol side chain reductase 2 (*SSR2*) by transcription activator-like effector nucleases (TALEN) suppressed the accumulation of toxic steroidal glycoalkaloids.^21^ Therefore, genome sequencing and gene annotation could facilitate the identification of target genes to enhance desired traits, such as higher contents of valuable compounds and lower contents of unwanted compounds. Additionally, the genome of edited plants could be compared with that of the reference genome to identify the potential risk of off-target changes.^22,23^ Long-read DNA sequencing technologies have emerged as powerful tools to obtain high-quality whole genome sequences.^24^ Recently developed PacBio technology has facilitated the generation of high-fidelity (HiFi) reads from circular consensus sequencing (CCS), with long-read (> 10 kb) and high accuracy (> 99%).^24^ In plant genome sequencing, highly contiguous near-chromosomal level sequences have been generated using HiFi read-only.^25^

Here, we generated a highly contiguous genome assembly of red perilla (*P. frutescens*) domesticated in Japan using PacBio HiFi reads. Functional annotation of identified genes was obtained using a systematic functional annotation workflow optimized for plants. It is anticipated that the highly contiguous genome assembly obtained in this study would promote the development of perilla varieties with desirable traits.

## 2. Materials and methods

### 2.1. Sample preparation and genome sequencing

DNA sample for genome sequencing was isolated from the young leaves of hydroponically grown *P. frutescens* cv. Hoko-3 using a Genomic-tip kit (Qiagen, Hilden, Germany). Library preparation was performed using SMRTbell Express Template Prep Kit 2.0 (PacBio, Menlo Park, CA, USA), and reads longer than 20 kb were collected using BluePippin (Sage Science, Beverly, MA, USA). Thereafter, the libraries were sequenced using Sequel IIe instrument (PacBio), and HiFi reads were selected from the circular consensus reads generated.

### 2.2. Omni-C sequencing

Dovetail Omni-C libraries were prepared using Omni-C kit (Dovetail Genomics, CA, USA), according to the manufacturer’s protocol. Sequencing was performed using DNBSEQ-G400 (MGI Tech, Shenzhen, China) in a 2 × 150 bp paired-end (PE) setting to obtain 356.34 million PE reads. The obtained reads were processed using fastp v0.23.1 software^26^ with default settings.

### 2.3. *De novo* assembly of HiFi reads

HiFi reads were assembled using Hifiasm v0.16.1^27,28^ with the combination of processed Omni-C reads with --hom-cov 55 --primary options (Hi-C integrated assembly).

### 2.4. Quality assessment of genome assemblies

Assembly statistics were obtained using QUAST v5.0.2^29^. Genome completeness was evaluated using BUSCO v5.2.2 software^30^ against embryophyta_odb10 (eukaryota, 2020-09-10) dataset (1,614 total BUSCO groups). *K*-mer based assembly evaluation was performed using Merqury v1.3^31^ and the Meryl db (k = 21) generated from the HiFi reads.

### 2.5. Omni-C scaffolding and construction of pseudochromosomes

The processed Omni-C reads were mapped to the primary contigs generated by Hifiasm using BWA-MEM v0.7.17,^32^ followed by the removal of the 3’-side of the chimeric mapping and merging of the paired BAM files using perl scripts from Arima Genomics mapping_pipeline (https://github.com/ArimaGenomics/mapping_pipeline) and SAMtools v1.12.^33^ The processed BAM file was converted to BED format using BEDtools v2.30.0,^34^ then scaffolding was performed using SALSA2 v2.3^35^ with -e DNASE -p yes options. To draw a Hi-C contact map, the output of SALSA2 was converted to .hic file using convert.sh script equipped with SALSA2, then the generated .hic file was visualized using Juicebox v1.11.08^36^. A total of 25 scaffolds of 1 Mb or longer were extracted using SeqKit v2.0.0^37^ and aligned with the 20 pseudochromosomes of the PF40 genome (GenBank assembly accession GCA_019511825.2^38^) using nucmer sequence aligner with default settings and filtering of the delta alignment file with delta-filter command with -q -r options in MUMmer4 v4.0.0rc1 package.^39^ The alignment was drawn with a custom R script (original: https://jmonlong.github.io/Hippocamplus/2017/09/19/mummerplots-with-ggplot2/). To create pseudochromosomes based on the alignment with the PF40 genome, scaffold_19 and the reverse complement sequences of scaffold_22 and scaffold_21 were joined in this order with gaps (500 Ns) to create a new scaffold (scaffold_19). The longest 20 scaffolds (scaffold_1–20) of this assembly were renamed following the PF40 genome. Additionally, 10 of the 20 scaffolds (scaffold_2, 3, 6, 8, 13, 14, 15, 16, 17, and 18) were reverse complemented to align with the PF40 genome in the same direction.

### 2.6. Removal of organellar sequences

The scaffolds were searched against chloroplast and mitochondrial genome sequences obtained from NCBI RefSeq (all genomic sequences available at mitochondrion and plastid directories, as of November 8, 2021) by blastn v2.12.0 with an E-value cutoff of 1e-5. Scaffolds with 90% or more coverage against one of the reference organellar sequences were labeled as organellar scaffolds and removed from the primary assembly. All of the scaffolds removed in this step were made from a single contig. We also performed the same organellar removal procedure for the alternate haplotigs generated by Hifiasm.

### 2.7. Transcriptome analysis

Total RNA for PacBio isoform sequencing (Iso-Seq) was extracted from the mixed sample of leaves, stems, and roots of the cultivar Hoko-3 using RNeasy Plant Mini kit (Qiagen, Hilden, Germany). Library preparation was performed using NEBNext Single Cell/Low Input cDNA Synthesis & Amplification Module (New England Biolabs, Ipswich, MA, USA) and SMRTbell Express Template Prep Kit 2.0 (PacBio). Thereafter, the reads were sequenced using the Sequel IIe instrument (PacBio), and CCS reads were generated using SMRTLink v10.0 (PacBio). The obtained BAM file was processed using IsoSeq v3.4.0 pipeline. Total RNA for RNA sequencing (RNA-Seq) was extracted from leaves of the cultivar Hoko-3 using ISOSPIN Plant RNA kit (Nippon Gene, Tokyo, Japan), and a sequencing library was prepared using NEBNext Poly(A) mRNA Magnetic Isolation Module (New England Biolabs) and NEBNext Ultra Directional RNA Library Prep Kit for Illumina (New England Biolabs). Sequencing was performed using NovaSeq 6000 (Illumina, San Diego, CA, USA) in a 2 × 150 bp paired-end (PE) setting. Reads from three biological replicates (154.20 million PE reads in total) were combined and used for gene prediction and annotation.

### 2.8. Gene prediction and annotation

The processed Iso-Seq reads (high-quality isoforms) were mapped to the assembled genome using minimap2 v2.23,^40^ then collapsed to obtain non-redundant isoforms using Cupcake ToFU scripts on cDNA_Cupcake v28.0.0 (https://github.com/Magdoll/cDNA_Cupcake). The RNA-Seq reads were processed using fastp v0.23.2 with default settings, mapped to the assembled genome using HISAT2 v2.2.1,^41^ and then transcript models were constructed using StringTie v2.2.1.^42^ Coding sequences from these two annotations were identified using GenomeTools v1.6.2,^43^ and gene features were obtained using a custom python script. The RNA-Seq reads were mapped to the assembled genome to predict protein-coding genes using BRAKER2 v2.1.6.^44^ For the input of BRAKER2, the assembled genome was repeat masked using RepeatModeler v2.0.2 and RepeatMasker v4.1.2 in Dfam TE Tools Container v1.4 (https://github.com/Dfam-consortium/TETools). To merge the annotations from Iso-Seq and RNA-Seq, non-overlapping gene features from RNA-Seq with those of Iso-Seq were obtained using BEDtools v2.30.0 intersect (-s and -v options), then concatenated with the annotations from Iso-Seq after removal of incomplete gene features using a custom python script. Similarly, non-overlapping gene features from BRAKER2 against those of the combined Iso-Seq and RNA-Seq were merged to obtain the combined gene annotations of Iso-Seq, RNA-Seq, and BRAKER2. The combined gene annotations were used for further analysis including functional annotation.

### 2.9. Functional annotation

Functional annotation of the protein-coding genes on the primary assembly was performed using Fanflow4Plants, designed for the functional annotation of plant species based on Fanflow4Insects.^45^ In the functional annotation of these protein coding sequences, these sequences were searched by GGSERACH v36.3.8g in the FASTA package (https://fasta.bioch.virginia.edu/). Instead of *Caenorhabditis elegans* and *Drosophila melanogaster* which were used as references in Fanflow4Insects, functionally well-curated protein datasets of *Arabidopsis thaliana, Oryza sativa*, and *Solanum lycopersicum*, as well as UniProtKB/Swiss-Prot, *Homo sapiens*, and *Mus musculus* were used as a reference (Supplementary Table S1). The sequences were also searched by HMMSCAN in HMMER package v3.3.2 (http://hmmer.org/) against the hidden Markov model (HMM) profile libraries of Pfam database v35.0.^46^

## 3. Results

### 3.1. *De novo* assembly of red perilla cultivar Hoko-3

We performed *de novo* assembly of the red perilla cultivar Hoko-3 (Fig. 1) from 72.4 Gb (57.5× coverage) of PacBio HiFi reads using Hifiasm. Hi-C (Omni-C) integrated assembly of Hifiasm was performed by combining the Omni-C reads and “--primary” option to generate primary and alternate contigs, as well as fully phased haplotype 1 and 2 contigs. We specified the homozygous coverage in the parameter setting of Hifiasm (--hom-cov 55) because the default setting could misidentify the coverage threshold for homozygous reads. The Hifiasm outputs generated 317 primary contigs and 14,150 alternate contigs (Supplementary Table S2). *K*-mer evaluation using Merqury showed that fully phased haplotype 1 and 2 were almost identical (Supplementary Fig. S1); therefore, we did not distinguish the fully phased haplotypes for further analysis. Merqury analysis indicated high base accuracy (QV = 60.1) and completeness (98.3%) of the primary contigs (Supplementary Table S2).

**Figure 1.**
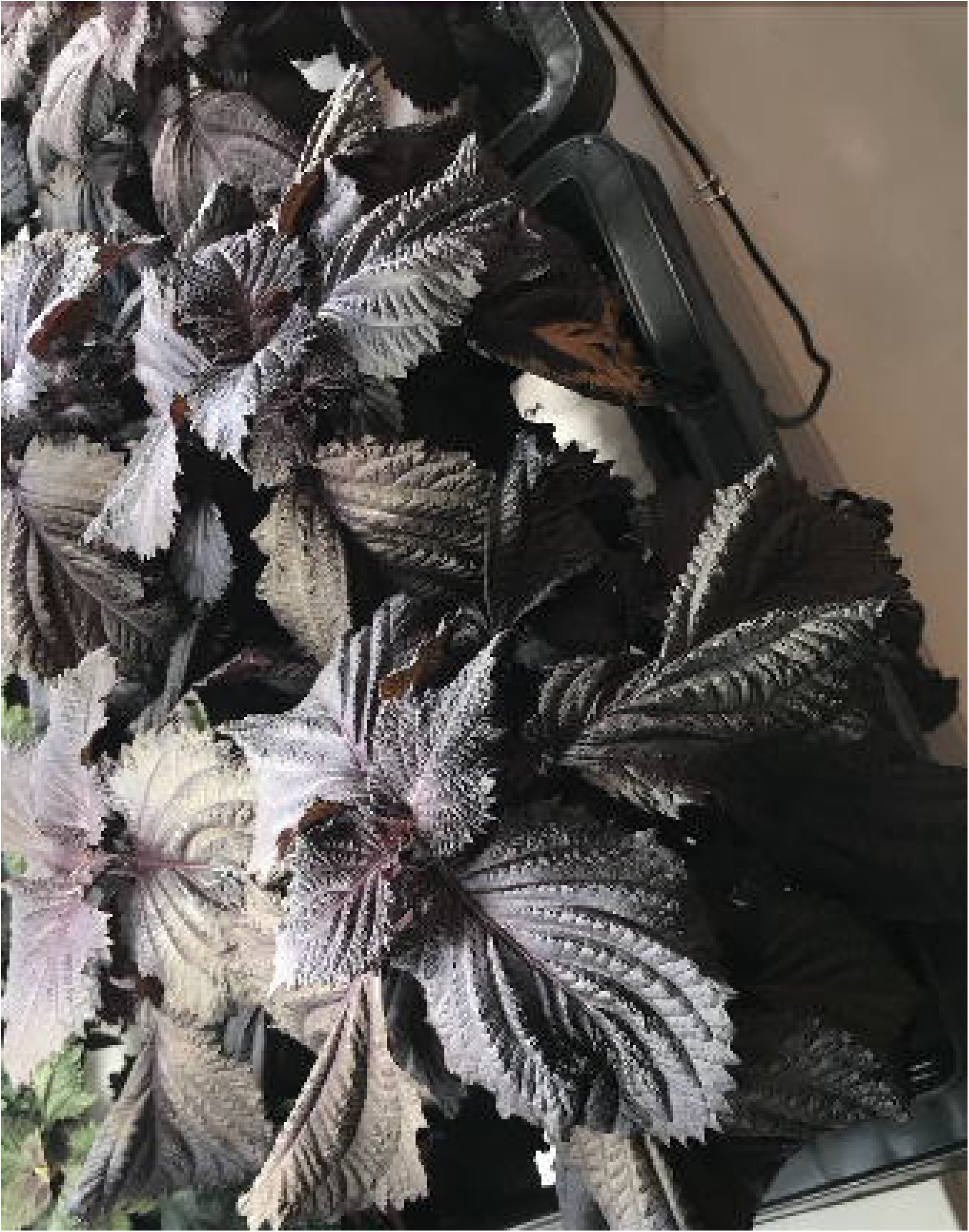
An image of the *Perilla frutescens* cv. Hoko-3 used for genome sequencing.

Additionally, the Omni-C reads were mapped to the primary contigs, and then scaffolding was performed using SALSA2 to construct pseudochromosomes. A total of 298 scaffolds were generated from the 319 primary contigs (two contigs were broken during the scaffolding), among which 25 were longer than 1 Mb (Supplementary Table S3). The 25 scaffolds (> 1 Mb) were aligned against the previously assembled chromosome-scale genome of green perilla cultivar PF40^38^ using MUMmer4 (Supplementary Fig. S2). Among the longest 20 scaffolds (scaffold_1–20), 19 (except for scaffold_19) covered each chromosome of the PF40 genome. Scaffold_19 partially covered chromosome 15 of the PF40 genome, and two other scaffolds (scaffold_21 and 22) partially aligned with chromosome 15 of the PF40 genome (Supplementary Fig. S2). Similarly, Hi-C contact map indicated that the three scaffolds (scaffold_19, 21, and 22) corresponded to the same chromosome (Supplementary Fig. S3); therefore, the three scaffolds were combined based on their partial alignment to chromosome 15 of the PF40 genome. We renamed and sorted the scaffolds based on the alignment with the PF40 genome to construct 20 pseudochromosomes (Fig. 2). After removal of scaffolds predicted to be derived from mitochondria or chloroplast genome, we obtained 71 scaffolds with N50 of 63.3 Mb (Pfru_yukari_1.0; Table 1). The N50 value of the scaffolds was almost similar to that of the PF40 assembly; however, the N50 value of the contigs was 41.5 Mb, which was ten times more than that of the PF40 assembly (Table 1). Each of the 20 pseudochromosomes consisted of less than six contigs, of which seven pseudochromosomes consisted of only one contig, indicating a highly contiguous genome assembly (Supplementary Table S4). Additionally, 99.2 % of the assembly was assigned to 20 pseudochromosomes (Supplementary Table S4). Completeness of the genome assembly evaluated with benchmarking universal single-copy orthologs (BUSCO) (Manni *et al.*, 2021) showed that the assembly achieved almost complete coverage of the BUSCO core gene sets (99.5% completeness) (Table 1). For the 14,150 alternate contigs produced by Hifiasm, we removed sequences predicted to be derived from mitochondria or chloroplast genome, and the obtained 12,627 contigs were deposited in DDBJ.

**Figure 2.**
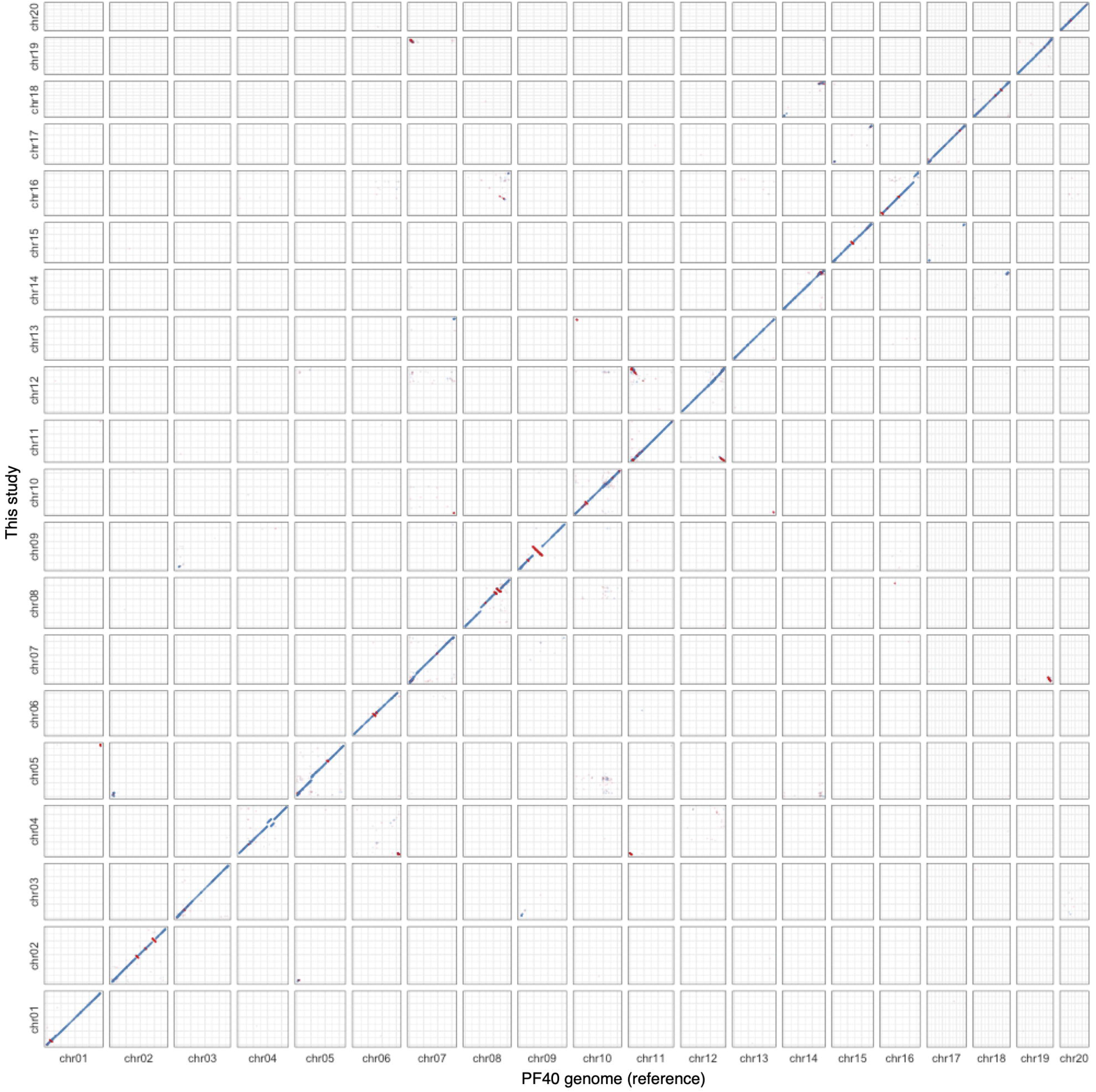
Dot-plot alignment between this genome assembly and reference PF40 genome assembly. Blue dots represent +/+ strand alignments and red dots represent +/- strand alignments. The large (> 3 Mb) structural differences detected in this alignment are eight inversions (two in chr02 and 08, and one in chr06, 09, 10, and 15), three gaps in PF40 genome (chr05, 08, and 16), and one rearrangement in chr04.

**Table 1.** Statistics of the genome assembly of red perilla cultivar Hoko-3 (Pfru_yukari_1.0) in comparison with the previous assembly of the PF40 genome (ICMM_Pfru_2.0)

|  | Pfru_yukari_1.0 | ICMM_Pfru_2.0 |
| --- | --- | --- |
| Total sequence length | 1,258,994,547 | 1,234,370,464 |
| No. of pseudochromosomes | 20 | 20 |
| No. of scaffolds | 71 | 1,465 |
| Scaffold N50 | 63,334,402 | 62,644,896 |
| Scaffold L50 | 9 | 10 |
| No. of contigs | 94 | 2,228 |
| Contig N50 | 41,459,669 | 2,738,655 |
| Contig L50 | 11 | 137 |
| Complete BUSCO<sup>a</sup> (%) | 99.5 | 99.3 |
| Complete and single-copy (%) | 4.1 | 6.3 |
| Complete and duplicated (%) | 95.4 | 93.0 |

<sup>a</sup>embryophyta\_odb10 (eukaryota, 2020-09-10) dataset (1,614 total BUSCO groups)

### 3.2. Annotation of the Hoko-3 genome

The length of repetitive sequences in the Hoko-3 genome was 866.7 Mb (68.84% of the genome) (Table 2). Long terminal repeat (LTR) elements accounted for 37.07% of the genome, with Copia and Gypsy constituting 14.01 and 14.92%, respectively (Table 2). Gene annotation of the Hoko-3 genome was performed by merging the gene models generated by Iso-Seq and RNA-Seq data, and gene prediction using BRAKER2 in this order. A total of 86,258 gene models were predicted with the 98.7% BUSCO complete data (Table 3). Iso-Seq, RNA-Seq, and BRAKER2 alone gave 66.7%, 97.0%, and 98.5% of BUSCO completeness, respectively (Supplementary Table S5). Although the Iso-Seq alone gave fewer number of genes (19,452 genes) and lower BUSCO completeness from the 33,869 transcripts (Supplementary Table S5), the combination of the Iso-Seq and RNA-Seq gave higher BUSCO completeness (97.7%), and addition of the predicted model from BRAKER2 achieved 98.7% BUSCO completeness (Table 3), suggesting that genes and isoforms identified in Iso-Seq contributed the higher completeness of the predicted gene models. The gene models were subjected to Fanflow4Plants, designed for the functional annotation of plant species based on Fanflow4Insects. Among the 86,258 gene models, 76,825 gene models were predicted as protein-coding genes, among which 72,983 gene models were annotated to at least one of the reference sequences in GGSERACH or pfam domain by HMMSCAN (Table 4).

**Table 2.**
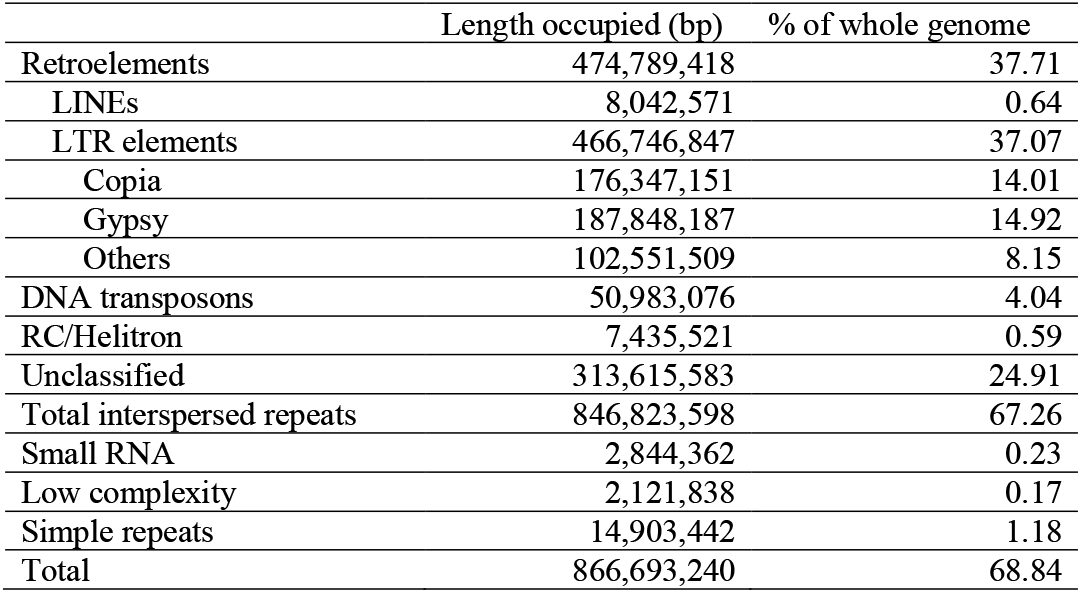
Summary of repetitive elements in Pfru_yukari_1.0.

|  | Length occupied (bp) | % of whole genome |
| --- | --- | --- |
| Retroelements | 474,789,418 | 37.71 |
| LINEs | 8,042,571 | 0.64 |
| LTR elements | 466,746,847 | 37.07 |
| Copia | 176,347,151 | 14.01 |
| Gypsy | 187,848,187 | 14.92 |
| Others | 102,551,509 | 8.15 |
| DNA transposons | 50,983,076 | 4.04 |
| RC/Helitron | 7,435,521 | 0.59 |
| Unclassified | 313,615,583 | 24.91 |
| Total interspersed repeats | 846,823,598 | 67.26 |
| Small RNA | 2,844,362 | 0.23 |
| Low complexity | 2,121,838 | 0.17 |
| Simple repeats | 14,903,442 | 1.18 |
| Total | 866,693,240 | 68.84 |

**Table 3.** Summary of the annotated genes.

|  | Iso-Seq | Iso-Seq<br>RNA-Seq | Iso-Seq<br>RNA-Seq<br>BRAKER2 |
| --- | --- | --- | --- |
| No. of genes | 19,452 | 53,541 | 86,258 |
| No. of transcripts | 33,869 | 88,890 | 121,996 |
| Complete BUSCO<sup>a</sup> | 66.7% | 97.7% | 98.7% |
| Complete and single-copy | 26.5% | 5.9% | 5.3% |
| Complete and duplicated | 40.2% | 91.8% | 93.4% |

<sup>a</sup>embryophyta\_odb10 (eukaryota, 2020-09-10) dataset (1,614 total BUSCO groups)

**Table 4.** Protein-level annotation of Pfru_yukari_1.0.

| Annotation Category | Annotation Level | Gene count |
| --- | --- | --- |
| Protein homolog from tophit | Arabidopsis | 54,262 |
|  | Rice | 55,694 |
|  | Tomato | 59,044 |
|  | Human | 38,255 |
|  | Mouse | 36,099 |
|  | UniProtKB/Swiss-Prot | 52,566 |
|  | At least one of the above | 72,339 |
| No protein homolog | Protein domain | 644 |
| Total genes with protein-level annotation |  | 72,983 |
| Hypothetical protein |  | 3,842 |
| Total |  | 76,825 |

### 3.3. Identification of the genes related to specialized metabolites in Hoko-3

As a practical example of genome editing target selection, enzyme coding genes in the anthocyanin biosynthetic pathway were studied. The major anthocyanin in *P. frutescens* is malonylshisonin, which is a glycosylated form of cyanidin.^3^ After the curation of the genes, the number of enzyme coding genes in the anthocyanin biosynthetic pathway in the genome were listed, including putative isoforms (Fig. 3, Supplementary Table S6). There were multiple copies of enzyme genes upstream of the pathway in the genome, but downstream enzyme genes were encoded in only a few locations in the genome. Based on this observation, it could be concluded that genes downstream of the pathway are most likely targets for genome editing to engineer the anthocyanin biosynthetic pathway in *P. frutescens*.

**Figure 3.**
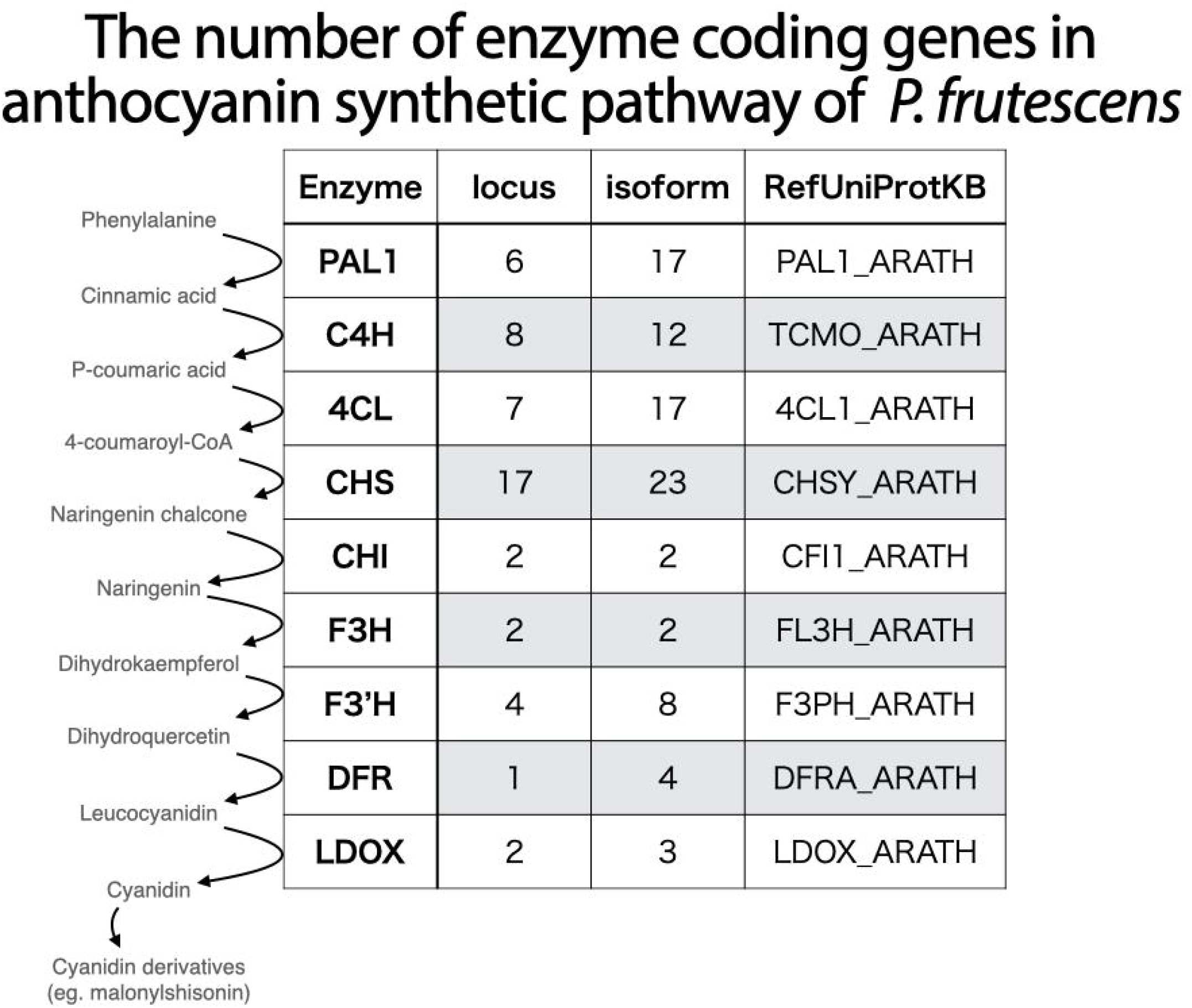
The number of enzyme coding genes in the representative anthocyanin biosynthetic pathway of *P. frutescens*. PAL, phenylalanine ammonia-lyase; C4H, trans-cinnamate 4-monooxygenase; 4CL, 4-coumarate:CoA ligase; CHS, chalcone synthase; CHI, chalcone-flavanone isomerase; F3H, flavanone 3-hydroxylase; F3’H, flavonoid 3’-hydroxylase; DFR, dihydroflavonol 4-reductase; LDOX, leucoanthocyanidin dioxygenase.

## 4. Discussion

Here, we generated a chromosome-level genome assembly of *P. frutescens* domesticated in Japan, using PacBio HiFi reads. Seven of the 20 pseudochromosomes were composed of only one contig, and the other pseudochromosomes were composed of not more than five contigs (Supplementary Table S4), indicating that the contigs generated from HiFi reads achieved a near-complete chromosome level. Recently, near-complete chromosome level assembly of *Macadamia jansenii* genome was generated from HiFi reads, with eight of the 14 pseudochromosomes represented by a single large contig,^25^ which is comparable to our assembly. Although it is difficult to construct complete chromosomal-level genome assembly from HiFi read-only, it is now possible to obtain near-complete chromosome-level assembly simply by running a HiFi read assembler, including Hifiasm. Additionally, in the scaffolding process using Omni-C reads, chromosome 15 was separated into three scaffolds. This is possibly due to some structural differences preventing these scaffolds from joining into a single sequence.

The number of gene models annotated in this study by combining two evidence-based annotations (Iso-Seq and RNA-Seq) and the gene prediction method (BRAKER2) was 86,258 (Table 3), which is almost twice the previously assembled *P. frutescens* genome (43,527 genes)^38^ and close to the number of genes reported in another Lamiaceae tetraploid species *Salvia splendens* (88,489 genes).^47^ The gene models generated in the present study achieved extremely high BUSCO completeness (98.7%) (Table 3), indicating that the models could be valuable resources for gene functional analysis of *P. frutescens*. Furthermore, an annotation system named Fanflow4Plants was developed based on the Fanflow4Insects for the functional annotation of gene models.^45^ Only well-curated protein datasets were used as references in this system to obtain reliable functional annotations, including protein sequences of three plant species (Arabidopsis, rice, and tomato) and two mammalian species (human and mouse), as well as UniProtKB/Swiss-Prot. Overall, 72,339 of 76,825 (94.2%) protein-coding genes were functionally annotated to at least one of the reference sequences (Table 4).

Since *P. frutescens* is a rich source of several metabolites,^1,8^ metabolic engineering could be used to enhance the biosynthesis and accumulation of valuable compounds in this species. Although genome editing of *P. frutescens* has not yet been reported, recent advances have shown that genome editing could be done using *Agrobacterium-mediated* transformation.^48^ In the present study, potential targets of genome editing to manipulate the anthocyanin biosynthetic pathway were identified (Fig. 3). As anthocyanin and rosmarinic acid share the upstream biosynthetic pathway towards 4-coumaroyl-CoA, it could be possible to change the metabolic flux into the biosynthesis of rosmarinic acid by knocking down the specific pathway for anthocyanin biosynthesis. Similar approaches could be used to identify target genes to enhance the biosynthesis of perillaldehyde or other beneficial compounds by examining the functional annotation of this genome assembly. Additionally, further analysis showed that the *P. frutescens* cv. Hoko-3 possessed a highly homozygous genome (Supplementary Fig. S1), which could be due to the fact that *P. frutescens* is a self-fertilizing crop.^49^ This homozygosity would be beneficial for the selection of unique targets for genome editing. Overall, our genome assembly and annotation could serve as a unique resource for future genome editing studies of *P. frutescens*.

## Acknowledgements

We thank Masaki Kurao (Hiroshima Prefectural Technology Research Institute) for technical assistance. This work was supported by Hiroshima Prefectural Government, the Center of Innovation for Bio-Digital Transformation (BioDX), an open innovation platform for industry-academia co-creation of JST (COI-NEXT, JPMJPF2010) and JSPS KAKENHI Grant 21K19118 to HB. Computations were partially performed on the NIG supercomputer at ROIS National Institute of Genetics.

## Author contributions

Conceptualization: KO, HB, YN

Methodology: KT, YT, MS, TM

Software: KT, YT, MS, HB

Validation: KT, YT, MS

Formal analysis: KT, YT, MS, HB

Investigation: KT, YT, MS, HB, SM

Resources: KT, YT, MS, HB, SM, YK, TI

Data Curation: KT, YT, MS, HB

Writing - Original Draft: KT

Writing - Review & Editing: MS, YT, TM, SM, YK, TI, KO, YN, HB

Visualization: KT, YT, MS, HB

Supervision: YN, HB

Project administration: HB

Funding acquisition: KO, HB

## Data availability statement

All sequencing data (assembled sequences and raw sequence reads) have been deposited in DDBJ under umbrella BioProject accession number PRJDB14288. The genome assembly from primary contigs has been deposited in DDBJ under the accession numbers BRKX01000001 to BRKX01000071. A set of haplotigs only sequences have been deposited in DDBJ under the accession numbers BRKY01000001 to BRKY01012627. The raw sequence reads have been deposited in DDBJ under the accession numbers DRR361636 (PacBio HiFi reads), DRR361637 (PacBio Iso-Seq reads), DRR361638 (Illumina RNA-Seq reads), and DRR415374 (Omni-C reads). Gene annotation and functional annotation of the protein coding genes are available at figshare (https://doi.org/10.6084/m9.figshare.20780995). Custom scripts used in this study are available at figshare (https://doi.org/10.6084/m9.figshare.20781466).

## Supplementary data

Supplementary data are available at figshare (https://doi.org/10.6084/m9.figshare.20780419).

## Legends for supplementary data

**Supplementary Table S1.** Source of protein-level functional annotation of Pfru_yukari_1.0.

**Supplementary Table S2.** Statistics of the contigs generated by Hifiasm.

**Supplementary Table S3.** AGP file (scaffolds_FINAL.agp) generated by SALSA2 describing the contig assignment of each scaffold.

**Supplementary Table S4.** Length and the components of 20 pseudochromosomes.

**Supplementary Table S5.** Summary of the gene annotation of each of the three methods.

**Supplementary Table S6.** The GGSERACH results corresponds to the identified genes in Figure 3.

**Supplementary Figure S1.** Merqury assembly spectrum plots of haplotype 1 (hap1) and haplotype 2 (hap2).

**Supplementary Figure S2.** Dot-plot alignment between the draft scaffolds (indicated as “s”) (longest 25 scaffolds; scaffold_1–25) and reference PF40 genome assembly. Blue dots represent +/+ strand alignments and red dots represent +/- strand alignments.

**Supplementary Figure S3.** Hi-C contact map of the draft scaffolds (indicated as “s”) generated by SALSA2.

## References

1. Ahmed, H. M. 2019, Ethnomedicinal, phytochemical and pharmacological investigations of *Perilla frutescens* (L.) Britt. Molecules, 24, 102.

2. Nitta, M., Lee, J. K., Kang, C. W., et al. 2005, The distribution of *Perilla* species. Genet. Resour. Crop Evol., 52, 797–804.

3. Saito, K., and Yamazaki, M. 2002, Biochemistry and molecular biology of the late-stage of biosynthesis of anthocyanin: lessons from *Perilla frutescens* as a model plant. New Phytol., 155, 9–23.

4. Ueda, H., Yamazaki, C., and Yamazaki, M. 2002, Luteolin as an anti-inflammatory and anti-allergic constituent of *Perilla frutescens*. Biol. Pharm. Bull., 25, 1197–202.

5. Deguchi, Y., and Ito, M. 2020, Rosmarinic acid in *Perilla frutescens* and perilla herb analyzed by HPLC. J. Nat. Med., 74, 341–52.

6. Longvah, T., Deosthale, Y. G., and Uday Kumar, P. 2000, Nutritional and short term toxicological evaluation of Perilla seed oil. Food Chem., 70, 13–6.

7. Hashimoto, M., Matsuzaki, K., Hossain, S., et al. 2021, Perilla seed oil enhances cognitive function and mental health in healthy elderly Japanese individuals by enhancing the biological antioxidant potential. Foods, 10, 1130.

8. Hou, T., Netala, V. R., Zhang, H., Xing, Y., Li, H., and Zhang, Z. 2022, *Perilla frutescens:* A rich source of pharmacological active compounds. Molecules, 27, 3578.

9. Yoshida, H., Nishikawa, T., Hikosaka, S., and Goto, E. 2021, Effects of nocturnal UV-B irradiation on growth, flowering, and phytochemical concentration in leaves of greenhouse-grown red perilla. Plants, 10, 1252.

10. Uemura, T., Yashiro, T., Oda, R., et al. 2018, Intestinal anti-inflammatory activity of perillaldehyde. J. Agric. Food Chem., 66, 3443–8.

11. Ji, W.-W., Wang, S.-Y., Ma, Z.-Q., et al. 2014, Effects of perillaldehyde on alternations in serum cytokines and depressive-like behavior in mice after lipopolysaccharide administration. Pharmacol. Biochem. Behav., 116, 1–8.

12. Sato, K., Krist, S., and Buchbauer, G. 2006, Antimicrobial effect of trans-cinnamaldehyde, (–)-perillaldehyde, (–)-citronellal, citral, eugenol and carvacrol on airborne microbes using an airwasher. Biol. Pharm. Bull., 29, 2292–4.

13. Tian, J., Wang, Y., Lu, Z., et al. 2016, Perillaldehyde, a promising antifungal agent used in food preservation, triggers apoptosis through a metacaspase-dependent pathway in *Aspergillus flavus. J. Agric*. Food Chem., 64, 7404–13.

14. Petersen, M., and Simmonds, M. S. J. 2003, Rosmarinic acid. Phytochemistry, 62, 121–5.

15. Yuba, A., Yazaki, K., Tabata, M., Honda, G., and Croteau, R. 1996, cDNA cloning, characterization, and functional expression of 4S-(–)-limonene synthase from *Perilla frutescens*. Arch. Biochem. Biophys., 332, 280–7.

16. Fujiwara, Y., and Ito, M. 2017, Molecular cloning and characterization of a *Perilla frutescens* cytochrome P450 enzyme that catalyzes the later steps of perillaldehyde biosynthesis. Phytochemistry, 134, 26–37.

17. Trócsányi, E., György, Z., and Zámboriné-Németh, É. 2020, New insights into rosmarinic acid biosynthesis based on molecular studies. Curr. Plant Biol., 23, 100162.

18. Nishida, K., and Kondo, A. 2021, CRISPR-derived genome editing technologies for metabolic engineering. Metab. Eng., 63, 141–7.

19. Nonaka, S., Arai, C., Takayama, M., Matsukura, C., and Ezura, H. 2017, Efficient increase of v-aminobutyric acid (GABA) content in tomato fruits by targeted mutagenesis. Sci. Rep., 7, 7057.

20. Ezura, H. 2022, Letter to the editor: The world’s first CRISPR tomato launched to a Japanese market: The social-economic impact of its implementation on crop genome editing. Plant Cell Physiol., 63, 731–3.

21. Sawai, S., Ohyama, K., Yasumoto, S., et al. 2014, Sterol side chain reductase 2 is a key enzyme in the biosynthesis of cholesterol, the common precursor of toxic steroidal glycoalkaloids in potato. Plant Cell, 26, 3763–74.

22. Graham, N., Patil, G. B., Bubeck, D. M., et al. 2020, Plant genome editing and the relevance of off-target changes. Plant Physiol., 183, 1453–71.

23. Sturme, M. H. J., van der Berg, J. P., Bouwman, L. M. S., et al. 2022, Occurrence and nature of off-target modifications by CRISPR-Cas genome editing in plants. ACS Agric. Sci. Technol., 2, 192–201.

24. Logsdon, G. A., Vollger, M. R., and Eichler, E. E. 2020, Long-read human genome sequencing and its applications. Nat. Rev. Genet., 21, 597–614.

25. Sharma, P., Masouleh, A. K., Topp, B., Furtado, A., and Henry, R. J. 2022, *De novo* chromosome level assembly of a plant genome from long read sequence data. Plant J., 109, 727–36.

26. Chen, S., Zhou, Y., Chen, Y., and Gu, J. 2018, fastp: an ultra-fast all-in-one FASTQ preprocessor. Bioinformatics, 34, i884–90.

27. Cheng, H., Concepcion, G. T., Feng, X., Zhang, H., and Li, H. 2021, Haplotype-resolved *de novo* assembly using phased assembly graphs with hifiasm. Nat. Methods, 18, 170–5.

28. Cheng, H., Jarvis, E. D., Fedrigo, O., et al. 2022, Haplotype-resolved assembly of diploid genomes without parental data. Nat. Biotechnol., 1332–5.

29. Mikheenko, A., Prjibelski, A., Saveliev, V., Antipov, D., and Gurevich, A. 2018, Versatile genome assembly evaluation with QUAST-LG. Bioinformatics, 34, i142–50.

30. Manni, M., Berkeley, M. R., Seppey, M., Simão, F. A., and Zdobnov, E. M. 2021, BUSCO Update: Novel and streamlined workflows along with broader and deeper phylogenetic coverage for scoring of eukaryotic, prokaryotic, and viral genomes. Mol. Biol. Evol., 38, 4647–54.

31. Rhie, A., Walenz, B. P., Koren, S., and Phillippy, A. M. 2020, Merqury: reference-free quality, completeness, and phasing assessment for genome assemblies. Genome Biol., 21, 245.

32. Li, H. 2013, Aligning sequence reads, clone sequences and assembly contigs with BWA-MEM. arXiv, 1303.3997.

33. Li, H., Handsaker, B., Wysoker, A., et al. 2009, The sequence alignment/map format and SAMtools. Bioinformatics, 25, 2078–9.

34. Quinlan, A. R., and Hall, I. M. 2010, BEDTools: a flexible suite of utilities for comparing genomic features. Bioinformatics, 26, 841–2.

35. Ghurye, J., Rhie, A., Walenz, B. P., et al. 2019, Integrating Hi-C links with assembly graphs for chromosome-scale assembly. PLoS Comput. Biol., 15, e1007273.

36. Durand, N. C., Robinson, J. T., Shamim, M. S., et al. 2016, Juicebox provides a visualization system for Hi-C contact maps with unlimited zoom. Cell Syst., 3, 99–101.

37. Shen, W., Le, S., Li, Y., and Hu, F. 2016, SeqKit: A cross-platform and ultrafast toolkit for FASTA/Q file manipulation. PLoS One, 11, e0163962.

38. Zhang, Y., Shen, Q., Leng, L., et al. 2021, Incipient diploidization of the medicinal plant *Perilla* within 10,000 years. Nat. Commun., 12, 5508.

39. Marçais, G., Delcher, A. L., Phillippy, A. M., Coston, R., Salzberg, S. L., and Zimin, A. 2018, MUMmer4: A fast and versatile genome alignment system. PLoS Comput. Biol., 14, e1005944.

40. Li, H. 2018, Minimap2: pairwise alignment for nucleotide sequences. Bioinformatics, 34, 3094–100.

41. Kim, D., Paggi, J. M., Park, C., Bennett, C., and Salzberg, S. L. 2019, Graph-based genome alignment and genotyping with HISAT2 and HISAT-genotype. Nat. Biotechnol., 37, 907–15.

42. Pertea, M., Pertea, G. M., Antonescu, C. M., Chang, T.-C., Mendell, J. T., and Salzberg, S. L. 2015, StringTie enables improved reconstruction of a transcriptome from RNA-seq reads. Nat. Biotechnol., 33, 290–5.

43. Gremme, G., Steinbiss, S., and Kurtz, S. 2013, GenomeTools: A comprehensive software library for efficient processing of structured genome annotations. IEEE/ACM Trans. Comput. Biol. Bioinform., 10, 645–56.

44. Brůna, T., Hoff, K. J., Lomsadze, A., Stanke, M., and Borodovsky, M. 2021, BRAKER2: automatic eukaryotic genome annotation with GeneMark-EP+ and AUGUSTUS supported by a protein database. NAR Genomics Bioinforma., 3, lqaa108.

45. Bono, H., Sakamoto, T., Kasukawa, T., and Tabunoki, H. 2022, Systematic functional annotation workflow for insects. Insects, 13, 586.

46. Mistry, J., Chuguransky, S., Williams, L., et al. 2021, Pfam: The protein families database in 2021. Nucleic Acids Res., 49, D412–9.

47. Jia, K.-H., Liu, H., Zhang, R.-G., et al. 2021, Chromosome-scale assembly and evolution of the tetraploid *Salvia splendens* (Lamiaceae) genome. Hortic. Res., 8, 177.

48. Kim, K.-H., Lee, Y.-H., Kim, D., et al. 2004, *Agrobacterium-mediated* genetic transformation of *Perilla frutescens*. Plant Cell Rep., 23, 386–90.

49. Sa, K. J., Kim, J. A., and Lee, J. K. 2012, Comparison of seed characteristics between the cultivated and the weedy types of *Perilla* species. Hortic. Environ. Biotechnol., 53, 310–5.

